# Mind the Gap: Identifying What Is Missed When Searching Only the Broad Scope with PubMed Clinical Queries

**DOI:** 10.1101/423111

**Authors:** Edwin V. Sperr

## Abstract

**Objective:** PubMed Clinical Queries are subdivided into “Broad” and “Narrow” versions. These versions are tuned to maximize either retrieval or sensitivity using two different sets of keywords and MeSH headings. While a searcher might assume that all items retrieved by *Filter name*/Narrow would also be found in the set *Filter name*/Broad, this is not explicitly guaranteed. It is the purpose of this study to quantify the overlap between these two sets and confirm whether *Filter*/Narrow is always a subset of *Filter*/Broad.

**Methods:** For each of the five PubMed Clinical Queries, PubMed was searched for citations matching the query *Filter*/Narrow NOT *Filter*/Broad. This number was compared with that for *Filter*/Broad to compute a “degree of discordance” between the two sets. This process was then repeated for the MeSH headings for “Medicine” and “Diseases” as well as for a set of test searches.

**Results:** Four of the five Clinical Queries returned citations using *Filter*/Narrow that were not found with *Filter*/Broad. Discordances between the sets Broad and Narrow were generally modest for “Etiology”, “Diagnosis” and “Clinical prediction guides”. “Prognosis” was notably more discordant – a searcher could easily miss one Prognosis/Narrow citation for every ten citations she retrieves when using Prognosis/Broad alone for a given search.

**Conclusions:** Users of the Clinical Queries apart from “Therapy” who are interested in retrieving as many relevant citations as possible should consider combining Filter/Narrow together with Filter/Broad in their search strategy. This is particularly true for “Prognosis”, as otherwise the risk of missing relevant citations is substantial.

## Introduction

It is widely appreciated that the rapid proliferation of the biomedical literature has made it difficult for health-care practitioners to keep pace. This is particularly true for full-time clinicians, who have little time to devote to locating resources. Many measures have been designed to deal with this issue, including the use of search “hedges”.

Search hedges (or “filters”) are strings of keywords and subject headings designed to combine with a user’s search to limit retrieval in a database. By ensuring that all results contain the words or concepts in the hedge, it is possible to increase the number of relevant results for a given search while simultaneously decreasing the effort required on the part of the searcher. Of course, it is difficult to imagine that end-users would use these tools if they were not already aware of them, so it is helpful that the National Library of Medicine has linked a tool called the “PubMed Clinical Queries” from the front page of PubMed (1).

The current form of the PubMed Clinical Queries is based on the work of R.B. Haynes and his group of researchers at McMaster University, who first published their work in 1994 (2). Over the following years, they further refined their work on the filters for “therapy” (3), “diagnosis” (4), “etiology” (5) “prognosis” (6) and “clinical prediction guides” (7). The development process for each filter involved compiling candidate text words and Medical Subject headings and then using those to develop test strategies that could be verified against a database of hand-selected “high quality” articles. The resulting strategies were then further refined and fine-tuned for sensitivity (the proportion of high quality articles retrieved) and specificity (how well the filter rejects low quality materials).

Variants of each filter were developed to emphasize different qualities, and two of those variants for each topic have been incorporated into PubMed Clinical Queries: Broad and Narrow. “Broad” queries are tuned to emphasize sensitivity and are intended for “those interested in comprehensive retrievals or in searching for clinical topics with few citations” (4). By contrast, the “Narrow” versions are optimized for specificity – “retrieval with little non-relevant material”. Table 1 shows how this is implemented for the concept of Prognosis (8).

**Table 1:** Clinical Query for “Prognosis”.

|  |  |
| --- | --- |
| **sensitive/broad**<br>**90%/80%** | (incidence[MeSH:noexp] OR mortality[MeSH Terms]<br>OR follow up studies[MeSH:noexp] OR prognos*[Text<br>Word] OR predict*[Text Word] OR course*[Text Word]) |
| **specific/narrow**<br>**52%/94%** | (prognos*[Title/Abstract] OR (first[Title/Abstract] AND<br>episode[Title/Abstract]) OR cohort[Title/Abstract]) |

Crucially, the Broad and Narrow versions of each query are not explicitly linked by terminology. Instead, each version was developed, tested and refined separately. This means, for example, that there is no guarantee that all items within the set *Filter name*/Narrow would also be within the set *Filter name*/Broad. It is logical that they would be – both sets of citations are relevant, and a user wanting to see as much useful material as possible would by necessity want to see both. It is the purpose of this study to quantify the overlap between these two sets and confirm whether *Filter*/Narrow is always a subset of *Filter*/Broad.

## Methods

To test whether the set *Filter*/Narrow is a subset of *Filter*/Broad, one can start with a simple search of PubMed using the fielded syntax: *Filter*/narrow[filter] NOT *Filter*/broad[filter]. If the number returned by this search is greater than zero, then some citations for that filter are being selected by Narrow while also being rejected by Broad.

If *Filter*/Narrow is not a subset of *Filter*/Broad, we can define a “degree of discordance” by calculating the following ratio: 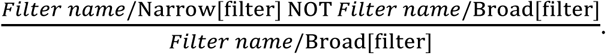 This quantities the significance of a discordance for a filter user employing *Filter*/Broad alone. A ratio of 0.001 means that for every thousand citations retrieved by Broad, one Narrow citation is missed. A ratio of 0.1 would mean that one Narrow citation is missed for every ten Broad citations retrieved.

While the focus of PubMed is human clinical medicine, it also includes basic science materials, veterinary science citations, etc., so we cannot make assumptions about the real-world performance of the Clinical Queries based solely of how well they perform in PubMed as a whole. It is straightforward to isolate and examine a subset of medicine-related citations by combining the above test with a search for “Medicine”[MeSH Terms]. As It is a reasonable assumption that Clinical Query users will employ them while researching a particular disease, this process should be repeated using the search “Diseases category”[MeSH terms]. Note, as the default behavior in PubMed is for subject heading searches to also include those other subjects indexed under them in the tree structure, these searches incorporate specific branches of medicine and individual diseases as well.

It is also important to note that it is entirely possible that discordances observed in large aggregates of citations might be different from those seen in the context of an individual search. To model real-world conditions, a list of sample searches was created and the discordance between Broad and Narrow sets was tested for each one. A list of 209 simple searches (9) were derived from a publicly-available list of common ICD-10 codes (10). These test searches were specifically chosen with an eye towards retrieving very large citation sets (“heart disease”) as well as smaller, more specific ones (“Vitamin D Deficiency Anemia”). A small Python program (11) was written to iterate through each query and test it programmatically against PubMed using the NCBI E-utilities API (12). The data resulting from the test searches was analyzed using Excel (Microsoft).

## Results

### Discordances for PubMed-in-total

PubMed was searched manually and programmatically in August 2018. The Therapy Clinical Query performed as one might expect – Narrow was demonstrated to be a subset of Broad, as there were no citations in the set Therapy/Narrow NOT Therapy/Broad. This was not the case for the other filters however. In PubMed-in-total, Etiology/Narrow NOT Etiology/Broad returned a bit over 53,300 citations for a computed degree of discordance of 0.0086. Larger discordances still were observed for the filters Clinical prediction guides (0.0271) and Diagnosis (0.0370), while Prognosis showed the largest discordance of all (0.0766). Figure 1 shows these relationships plotted as proportional Venn diagrams (13).

**Figure 1:**
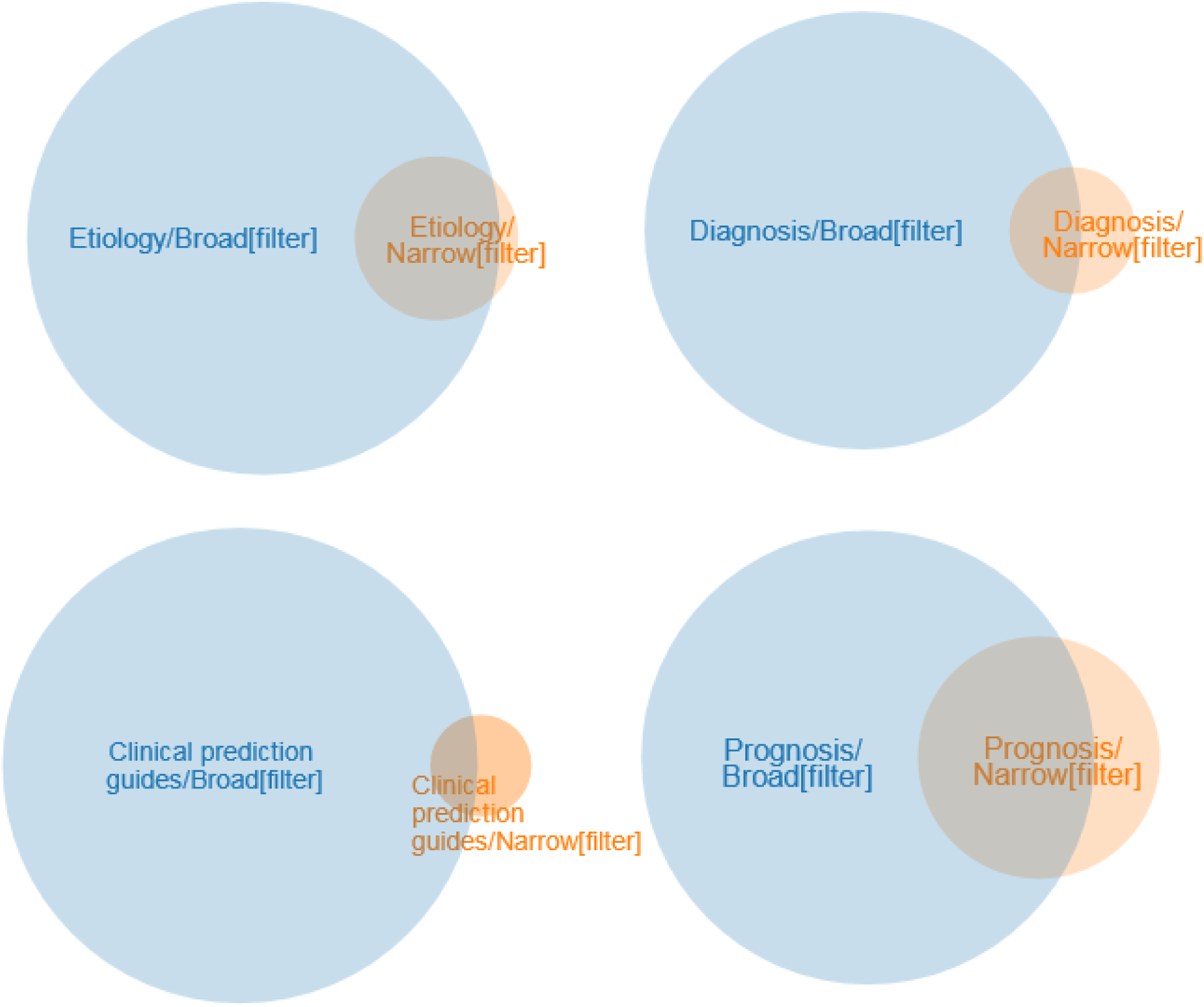
*Filter*/Narrow vs. *Filter*/Broad in PubMed.

### Discordances for “Diseases” and “Medicine”

A similar pattern is seen for the citations indexed under “Diseases category”[MeSH Terms] and “Medicine” [MeSH Terms]. Etiology again shows the smallest discordances between Narrow and Broad (and slightly smaller ratios for the MeSH subsets than for PubMed as a whole), with ratios of 0.0063 for Diseases and 0.0076 for Medicine. An examination of the citations selected by Clinical prediction guides shows ratios for “Diseases” (0.0196) and “Medicine” (0.0447) that bracket that for PubMed-in-total. “Medicine” (0.0086) and “Diseases” (0.0117) subsets of Diagnosis were notably less discordant than in PubMed-in-total. As above, Prognosis stands out as the most discordant of the four, with “Diseases” exhibiting a ratio of 0.0742 and “Medicine” one of 0.0811. (Table 2)

**Table 2:** Degrees of Discordance between *Filter*/Narrow and *Filter*/Broad.

| Filter | All PubMed | Diseases | Medicine | Test Queries |  |
| --- | --- | --- | --- | --- | --- |
|  |  |  |  | Mean | Median |
| Etiology | 0.0086 | 0.0063 | 0.0076 | 0.0088 | 0.0083 |
| Clinical prediction guides | 0.0271 | 0.0196 | 0.0447 | 0.0148 | 0.0142 |
| Diagnosis | 0.0370 | 0.0117 | 0.0086 | 0.0083 | 0.0055 |
| Prognosis | 0.0766 | 0.0742 | 0.0811 | 0.1076 | 0.0989 |

### Discordances for Test Searches

For the test query set, Diagnosis showed the least amount of discordance (mean: 0.0083, median: 0.0055, range: 0 - 0.0432, SD: 0.0078), while Etiology had similar values (mean: 0.0088, median: 0.0083, range: 0 - 0.0211, SD: 0.0040). Test queries for Clinical prediction guides were slightly more discordant (mean: 0.0148, median: 0.0142, range: 0.0021 - 0.0371, SD: 0.0068). Finally, degrees of discordance for Prognosis were again by far the highest of the four. Indeed, ratios for Prognosis test queries tended to be even higher than those for any of the Prognosis citation aggregates (mean: 0.1076, median: 0.0989, range: 0.0200 - 0.3824, SD: 0.0561). Data for all four sets of test queries are summarized in Figure 2.

**Figure 2:**
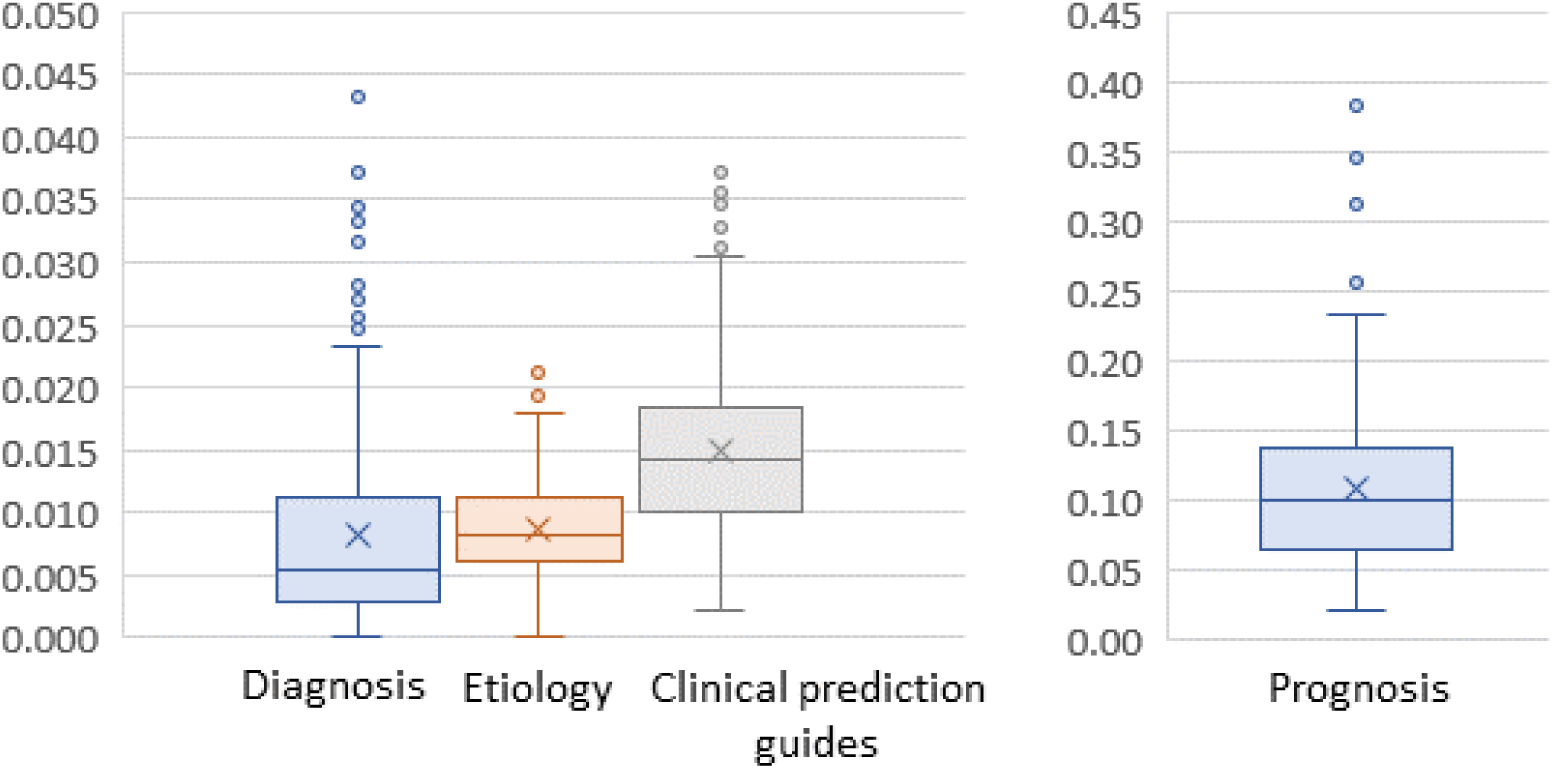
Degrees of Discordance for test searches.

### Differentiating Narrow NOT Broad citations

A comprehensive study of how citations in the sets *Filter*/Narrow NOT *Filter*/Broad differ from other Narrow citations is beyond the scope of this work, but we can attempt to illustrate some aspects of these aggregates. Just as one can locate discordant items via the search terms noted before, one can find those Narrow items that *are* contained within Broad by searching “*Filter name*/Narrow[filter] AND *Filter name*/Broad[filter]”. Once we have defined searches that can differentiate the two portions of Narrow, we can examine those citations by combining them both with other search terms that define a possibly-relevant aspect. (see Table 3)

**Table 3:** Proportion of citations within selected categories for each portion of *Filter*/Narrow.

| Category | Etiology |  | Clinical prediction guide |  | Diagnosis |  | Prognosis |  |
| --- | --- | --- | --- | --- | --- | --- | --- | --- |
|  | AND | NOT | AND | NOT | AND | NOT | AND | NOT |
| Diseases Category (subject) | 74.0% | 47.7% | 48.9% | 36.7% | 71.3% | 23.5% | 83.3% | 66.7% |
| Humans (subject) | 86.8% | 57.7% | 65.7% | 58.0% | 77.6% | 42.3% | 86.6% | 79.1% |
| Published from 2008 onwards | 65.8% | 81.2% | 75.6% | 70.7% | 55.3% | 35.7% | 59.1% | 74.4% |
| Comparative Study (pub. type) | 11.7% | 4.5% | 8.6% | 7.3% | 17.2% | 8.4% | 8.6% | 8.9% |
| Clinical Study (pub. type) | 7.3% | 5.9% | 5.0% | 2.5% | 5.3% | 0.8% | 5.9% | 7.4% |
| Core clinical journals subset | 13.7% | 7.2% | 6.5% | 3.7% | 10.2% | 4.5% | 11.5% | 11.4% |

For each filter, fewer Narrow NOT Broad citations were indexed with the subject headings “Diseases Category”[MeSH Terms] or “Humans”[MeSH Terms] than their Narrow AND Broad counterparts. Beyond that, it is difficult to generalize; some categories show sharp discrepancies for one or two filters, but not in others. It should be noted that Prognosis, the filter that consistently shows the largest degree of discordance between Broad and Narrow NOT Broad, has similar percentages within several categories.

## Discussion

It has been shown that there is a consistent pattern where some of the citations that get selected by the Narrow filters in four out of the five Clinical Queries are simultaneously rejected by their Broad counterparts. It has been further demonstrated that this effect appears consistently across different citation aggregates as well as in the types of searches that end users are likely to perform. This has clear implications for search effectiveness – a searcher using Broad to obtain “comprehensive retrievals” will be frustrated, as some relevant citations are nonetheless not being retrieved.

This effect varies in intensity; using Diagnosis/Broad is likely to mean missing at least five out of 1000 relevant citations, while with Etiology/Broad that number would probably climb to eight. It should be noted however that individual searches can show larger discordances. Many of the queries tested against Etiology/Broad missed one Narrow citation for every hundred Broad citations retrieved. A search using Diagnosis/Broad could easily miss twice that number. Of course, without manually checking, a user has no way of knowing where on this continuum her individual search might lie.

It is with the Prognosis filter that this effect becomes truly worrisome. Using Prognosis/Broad alone, a searcher is likely to miss seven or eight Narrow citations for every 100 Broad retrieved. Indeed, out of the 209 individual test queries searched, 103 had a ratio of higher than 0.1, while eleven had a ratio of higher than 0.2. The latter ratio demonstrates that, when using Prognosis/Broad alone, there is a non-trivial chance of missing two out of every ten relevant citations.

It is difficult to gauge the quality of Narrow NOT Broad citations when compared to other Narrow citations, but it possible to make some general observations. Compared to Narrow AND Broad citations, Narrow NOT Broad citations are generally less likely to be indexed with “Diseases category”[Mesh] or “Humans”[Mesh], which could be taken as an indicator of lesser clinical relevance. By contrast, Prognosis/Narrow NOT Prognosis/Broad citations are slightly more likely to be described with the publication types “Comparative Study” or “Clinical Study”, which could well be evidence of higher quality. Indeed, the clinical relevance of Prognosis/Narrow NOT Prognosis/Broad citations in particular is further demonstrated by the appearance of many of them in the hand-curated McMaster Plus database (14). Inclusion in that database is the primary test criteria used in the 2013 revalidation study of the Clinical Queries by Wilczynski, *et. al.* (15).

Perhaps more to the point, all *Filter*/Narrow NOT *Filter*/Broad citations are firstly in the set *Filter*/Narrow. They are selected by the same system of keywords and subject headings as other Narrow citations, so it would be difficult to dismiss the importance of their exclusion without calling the effectiveness of the entire hedge into question. This is especially true as substantial portions of all Prognosis/Narrow (27%), Diagnosis/Narrow (44%) and Clinical prediction guides/Narrow (56%) citations are indeed Narrow NOT Broad.

The data shown above demonstrate that PubMed users should be aware of the limitations of the Clinical Queries when using them to search. For Etiology, Diagnosis, Clinical Prediction Guides and Prognosis, the differences in retrieval between Filter/Broad and Filter/Narrow NOT Filter/Broad mean that there are sometimes many relevant citations that are missed when searching with Broad alone. The importance of these missing citations is to some degree dependent on the task at hand. If one is merely attempting to find a couple of reviews of a common condition using Diagnosis/Broad, it might make little difference if one or two citations out of 100 are missed. If one is constructing a literature review however, the lack of five citations out of 1000 could have a significant impact.

In short, users of the Clinical Queries for Etiology, Diagnosis and Clinical Prediction Guides who are interested in retrieving as many relevant citations as possible should consider the expedient of combining *Filter*/Broad with Filter/Narrow with a Boolean OR in their search strategy. This is even more the case when using Prognosis – otherwise the risk of missing relevant citations is substantial.

